# Safety and Tolerability of Bacteriophage Therapy in Severe *Staphylococcus aureus* Infection

**DOI:** 10.1101/619999

**Authors:** Aleksandra Petrovic Fabijan, Ruby CY Lin, Josephine Ho, Susan Maddocks, Jonathan R Iredell, on behalf of the Westmead Bacteriophage Therapy Team (WBTT), AmpliPhi Biosciences Corporation

## Abstract

**Importance:** The effect of IV administration of a bacteriophage cocktail produced under GMP conditions on patients with severe *S. aureus* infection, including complicated bacteraemia, endocarditis and septic shock, is unknown.

**Objective:** To assess safety and tolerability of adjunctive bacteriophage therapy in patients with severe *S. aureus* infections.

**Design, Setting, Participants:** Observational, open-label clinical trial of thirteen critically-ill patients admitted to a tertiary-referral hospital with *S. aureus* bacteraemia (including infective endocarditis, n=6) were assessed by the treating clinician and two consulting infectious diseases physicians to independently verify that routine medical and surgical therapy was optimal and that a poor outcome remained likely. Compassionate access to therapy was approved by both US and Australian regulators and by the Westmead Hospital Human Research Ethics Committee.

**Intervention:** A GMP-quality preparation of three combined *Myoviridae* bacteriophages with specific activity against *S. aureus* (AB-SA01), was administered intravenously in conjunction with optimal antibiotic therapy.

**Main Outcome and Measurements:** Physiological, haematological and biochemical markers of infection, bacterial and bacteriophage kinetics in blood, development of resistance to bacteriophages, and mortality at 28 (D28) and 90 (D90) days were measured. Main outcomes were safety and tolerability.

**Results:** Bacteriophage therapy was initiated 4-10 days after antibiotic commencement, at 10^9^ plaque-forming units (PFU) twice daily. Infecting staphylococci were typical of common local subtypes. Initial input ratio of phages to bacteria in the bloodstream (MOI_input_) was >100. Five of the thirteen patients died by D28 and a sixth patient suffered sudden cardiac death on D90. Bacteriophage therapy coincided with a marked reduction in staphylococcal bacterial DNA in the blood and in sepsis-associated inflammatory responses in almost all cases. No bacteriophage-attributable adverse events were identified. Development of bacteriophage resistance was not observed. Population analysis revealed no significant effect of bacteriophage therapy on the gut microflora.

**Conclusions and Relevance:** Adjunctive bacteriophage therapy appears to be safe and well-tolerated in critically ill patients with severe *S. aureus* infection. Two weeks of twice daily intravenous administration may be a suitable protocol. Controlled trials are needed.

**Trial Registration:** Westmead Hospital Human Research Ethics Committee approval July 11, 2017; ClinicalTrials.gov Identifier: NCT03395769, AB-SA01-EAP01 (January 10, 2018); Clinical Trials Notification (Australian Therapeutic Goods Association): CT-2018-CTN-02372-1 (July 23, 2018).

**Key Points:** 

**Question:** Is intravenous (IV) administration of investigational bacteriophage (phage) therapy safe and well-tolerated in patients with severe *Staphylococcus aureus* infection?

**Findings:** Thirteen patients with severe *S. aureus* infections received AB-SA01, a bacteriophage product prepared according to Good Manufacturing Practices (GMP), as adjunctive therapy to antibiotics. AB-SA01 was well-tolerated with no adverse events identified. Bacterial burden and inflammatory responses were reduced and no phage-resistant staphylococci were isolated during or after therapy.

**Meaning:** Our results will inform future randomised controlled trials assessing the antibacterial and anti-inflammatory potential of bacteriophages in the treatment of severe *S. aureus* infection.

## INTRODUCTION

*S. aureus* bacteraemia is associated with a 30-day mortality of ∼15%y. This nearly doubles in the setting of infective endocarditis (IE), of which it is the leading cause^1^, and attributable mortality is higher again in those with prosthetic valve infection^2^.

Bacteriophages (phages) are natural viral predators of bacteria that have been used therapeutically for over a century, including for *S. aureus* sepsis^3^. Long overshadowed by antibiotics, phage therapy is now experiencing a renaissance as adjunctive therapy to antibiotics and for severe antibiotic-resistant infections. AB-SA01 (AmpliPhi Biosciences) is a GMP-quality preparation of three *Myoviridae* bacteriophages (10^9^ PFU of each) with broad activity against *S. aureus*, the safety of which was recently demonstrated for intranasal therapy of chronic rhinosinusitis^4^. Here, we describe a single-centre experience with thirteen cases of intravenous AB-SA01 therapy for severe *S. aureus* infections including prosthetic valve endocarditis (PVE) and septic shock.

## METHODS

An open-label protocol was approved by our institution’s Ethics Committee for adjunctive therapy with AB-SA01. Informed consent was sought from patients with two or more consecutive daily blood cultures positive for *S. aureus* if the treating clinician and two independent infectious diseases specialists agreed that treatment was already optimal, significant risk of poor outcome remained and imminent demise (<48hours) was not expected (Figure 1). AB-SA01 was administered intravenously twice daily for 14 days in 50-100 mL of 0.9% sodium chloride over 10-30 minutes. An echocardiogram was performed and the presence of IE determined according to modified Duke criteria^5^. Vital signs were monitored during and immediately after phage administration for up to 4 hours after the first dose and clinical, haematological and blood biochemical parameters were monitored out to 90 days in a standardised protocol (Table 1). Charlson comorbidity indices (CCI, a guide to one-year mortality on the basis of pre-existing comorbidities)^6^, Acute Physiology and Chronic Health Evaluation (APACHE) II scores (a predictor of in-hospital mortality for intensive care cohorts)^7^ and IE mortality risk scores (a 6-month mortality predictor for IE)^8^ were calculated. *S. aureus* isolates were tested for susceptibility to antibiotics (BD Phoenix, USA) and to AB-SA01 and their DNA sequences were determined.

**Table 1:** Patient data

| WMP | Diagnoses | *S. aureus* ST | Predicted % mortality |  |  | Antibiotics and phage therapy |  |  | CRP<br>WBC |  | Outcome | Comments |
| --- | --- | --- | --- | --- | --- | --- | --- | --- | --- | --- | --- | --- |
|  |  |  | CCI | APACHE II (score) | IE | Principal antibiotics<sup>a</sup> | (days prior<sup>b</sup>) | Phage doses | Pre<sup>c</sup> | D14<sup>e</sup> |  |  |
| 001 65yM | Septic shock, PVE (A, M) | 15 | 23 | 25 (18) | 87 | FLU, CIP, RIF, MEM | (10) | 28 | 171<br>24.5 | 47<br>16.2 | Survived D28<br>Survived D90 | - |
| 003 47yM | Septic shock, IE (A, M) | 340 | 23 | 40 (25) | 68 | FLU | (5) | 5 | ND<br>17.9 | - | Died D3 | Septic shock; valve failure |
| 004 43yF | OM/epidural abscess | 72 | 2 | 8 (6) | 17 | FLU | (6) | 28 | 271<br>6.2 | 50<br>4.4 | Survived D28<br>Survived D90 | - |
| 005 39yM | OM/epidural abscess | 25 | 2 | 4 (3) | 10 | FLU | (6) | 28 | 19<sup>d</sup><br>13 | 28<br>7.1 | Survived D28<br>Survived D90 | - |
| 006 87yM | Possible PVE (A) | 5* | 98 | 40 (23) | 74 | FLU | (8) | 4 (w) | 151<br>26.3 | - | Died D6 (w) | Refused CRRT |
| 007 89yM | OM/septic arthritis | 109 | 98 | 40 (21) | 35 | FLU | (7) | 24 (w) | 130<br>5.3 | - | Died D15 (w) | Myelodysplastic syndrome, palliated D13 |
| 008 36yF | RIE (T), empyema | 15 & 398* | 2 | 8 (7) | 26 | FLU, CLI | (7) | 28 | 252<sup>d</sup><br>13.9 | ND<br>9.6 | Survived D28<br>Survived D90 | 2 months later: ST672 sepsis |
| 009 69yF | PVE (A, M) | 45 | 79 | 40 (25) | 45 | FLU, CIP, RIF | (8) | 28 | 149<br>45.8 | 48<br>9.4 | Survived D28<br>Died D90 | Surgery D50 |
| 010 21yM | PVE (A) | 5 | 2 | 25 (17) | 40 | FLU, CIP, RIF | (6) | 28x2 | 79<br>16.1 | 241<br>22.9 | Survived D28<br>Survived D90 | Surgery D15; phage x2 courses |
| 011 64yM | Septic shock, pneumonia | (22)* | 10 | 40 (25) | 45 | VAN, CLI, MEM, AZI | (5) | 5 (w) | 437<sup>d</sup><br>14.9 | - | Died D7 (w) | ECMO, CRRT |
| 012 71yM | Epidural abscess | 93 | 23 | 15 (12) | 35 | CEF, CLI, FLU | (6) | 28 | 164<br>9 | 15<br>7.4 | Survived D28<br>Survived D90 | - |
| 013 81yM | PVE (A) | 45* | 100 | 25 (19) | 50 | FLU, CIP, RIF | (4) | 27 | 156<br>15.2 | 96<br>12.7 | Died D27 | Respiratory failure |
| 014 70yM | Possible IE (A) | 188 | 47 | 8 (6) | 40 | CEF, FLU | (4) | 28 | 327<sup>d</sup><br>6.9 | 19<br>5.2 | Survived D28<br>Survived D90 | Mycotic aneurysm |

**Figure 1.**
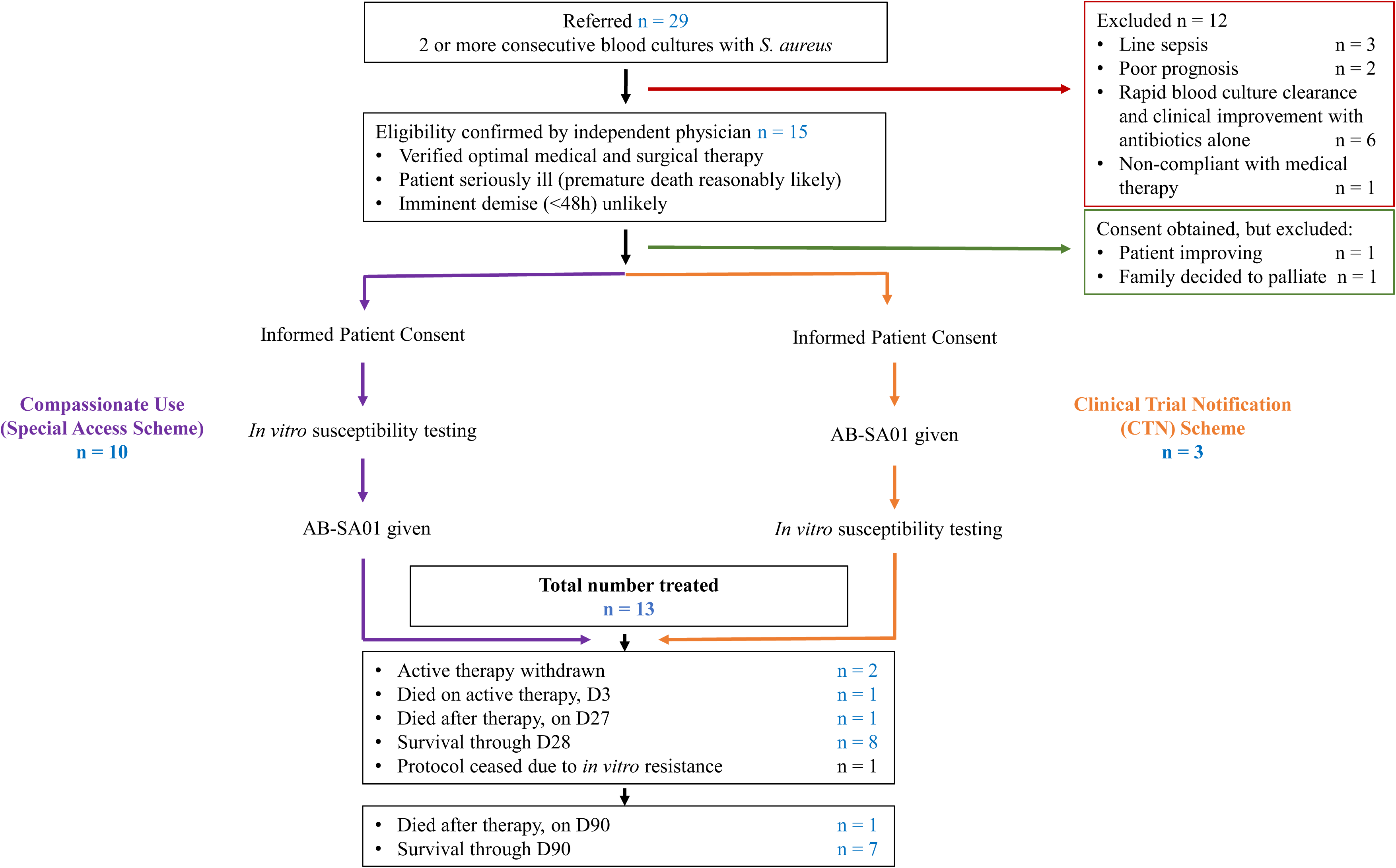
Study outline from consent through to D90 follow-up. Individuals with severe *S. aureus* infections accessed AB-SA01 through TGA’s compassionate use Special Access Scheme (SAS). Patient consented after 23^rd^ of July 2018 were under TGA’s Clinical Trial Notification (CTN) scheme. All cases were notified to the US FDA and the Australian TGA.

Quantitative PCR targeting *nuc* and *femA* of *S. aureus* and the primase gene of *Myoviridae* (5’-AGAGAGAGCCCCTTTACTTAC - 3’ forward and 5’- CCCTTCAGGTAATCGAGGAGGA - 3’ reverse) was performed on DNA extracted from whole blood (*S. aureus*) and serum (phage) respectively (Figure 2). Viable phages were quantified in serum by diluting onto lawns of a flucloxacillin-, ciprofloxacin-and rifampicin-resistant MRSA strain (SA-5863) of multi-locus sequence type 239, which yielded clear plaques by a standard method^9^ at ten-fold dilutions of the AB-SA01. Phage efficiency of plating (EOP, an indirect *in vitro* measure of lysis efficiency) was determined for each isolate in triplicate. RNA was extracted from whole blood^10^ for transcriptomic analysis (Figure 2C). Bacterial population diversity was assessed by *16S rRNA* gene amplicon sequencing of DNA from faecal samples^11^.

**Figure 2.**
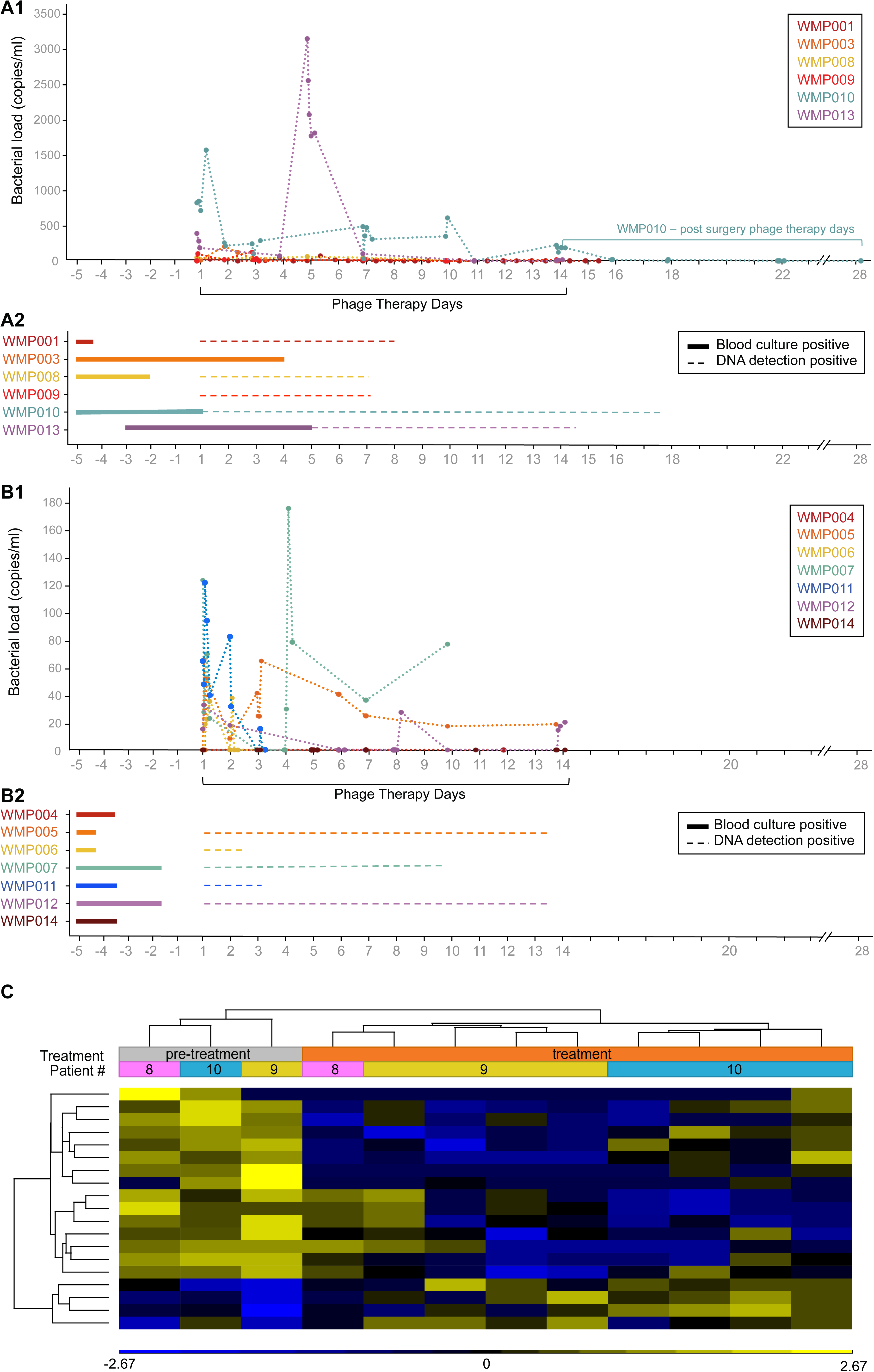
*S. aureus* DNA burden in the blood: **A)** endocarditis patients and **B)** bacteremia patients, and their corresponding blood culture positivity before and during phage therapy for endocarditis 2A1 and bacteriemia, 2B1. DNA was extracted from EDTA, Heparin (Patient 1-5) and PAXgene blood tubes (Patient 6-13) using Qiagen Blood DNA Mini kit according to manufacturer’s instructions. Bacterial load was then determined by multiplex-tandem polymerase chain reaction with the AusDiagnostics STAPHYLOCOCCUS + VRE (8-WELL) kit according to manufacturer’s instructions. Standard curves were generated with 1 mL of blood obtained from healthy individual and spiked with normal saline (0.9% NaCl) containing log-phase *S. aureus* in 10-fold serial dilutions. **C)** Transcriptome analysis of whole blood from endocarditis patients (WMP008, 9 and 10) using NanoString PanCancer Immune Profiling Panel (NanoString Technologies, Seattle, WA). Normalisation to housekeeping genes and QC checks were performed in nSolver (NanoString, v4.0) and then log 2 transformed (Partek Genome Suite v7.0). One-way ANOVA (p<0.05 unadjusted) indicates down-regulation of the innate immune response within 72h of phage therapy, with hierarchical clustering analysis revealing nineteen significantly differentially expressed genes, identified in Fig. 2C according to their KEGG functional description.

## RESULTS

Thirteen patients with severe complicated *S. aureus* bacteraemia received AB-SA01 from August 2017 to September 2018, inclusive (Figure 1 & Table 1). No fever, rash, hypotension or any other adverse events were reported during or after AB-SA01 infusion. All received flucloxacillin (n=10), cefazolin (n=2) or vancomycin (n=1) as primary therapy for six weeks or until active therapy was ceased, often with additional ciprofloxacin and/or rifampicin (Table 1), with infectious diseases specialist supervision throughout. Phage therapy began within 4-10 days of antibiotics.

Three died in the first week (WMP003, WMP006 and WMP011), two after withdrawal of therapy due to expected poor outcomes and one (WMP003) during attempted surgical source control after only 48 hours of phage therapy following 8 days of unremitting bacteraemic shock on high-dose antibiotics (Table 1). All blood cultures and intraoperative specimens from WMP003 grew AB-SA01-susceptible *S. aureus* (a leukocidin *lukGH*-positive ST340 strain). All others showed improvement by day 14. In all, six deaths occurred in the 90 day follow-up, of which five (38%) were within 28 days of starting phage therapy, with an additional sudden death at home on D90 (WMP009) attributed to (known) ventricular dysrhythmia.

In all, two patients had possible IE and six patients had definite IE, of whom four had PVE (Table 1). Two PVE cases (WMP001 and WMP010) and WMP014 (possible IE) were complicated by clinically significant cerebral emboli. WMP010 received a second two-week course beginning on the day of prosthetic aortic valve replacement (for incompetence) after three weeks of high-dose antibiotics: resected paravalvar tissues were culture-negative but *S. aureus* and *Myoviridae* DNA was detected. WMP009 had metallic aortic and mitral valves replaced more than a month after finishing phage therapy when well and afebrile (on D58 of antibiotics; all resected tissues were culture negative) and WMP001 avoided valve surgery altogether.

*S. aureus* isolates were mostly (12/13) methicillin-susceptible and of common sequence types, including a single methicillin-resistant (MRSA) strain (Table 1). *S. aureus* isolates from WMP011 were not lysed at all by AB-SA01 phages *in vitro* and EOP was reduced in two others (WMP006 and WMP013) (Table 1). Patients infected with staphylococci in which AB-SA01 produced clear plaques at >10^-6^ dilutions *in vitro* (10/13), with a predicted 6 months mortality score ∼40%^5^, suffered an early mortality of 20% (2/10 at 28 days) and total mortality of 30% (3/10 at 90 days).

Bacteriophage susceptibility *in vitro* did not change in any case during or after infection. Two unrelated bacteraemic strains of *S. aureus* (an AB-SA01-susceptible ST15 strain and a non-susceptible ST398 strain) were identified in one patient (WMP008) who was lost to follow-up before D90. She was readmitted two months later with sepsis after ongoing recreational self-injection and AB-SA01-susceptible *S. aureus* (ST672) was repeatedly cultured from blood with no evidence found of either previous strain.

Bacterial DNA load in blood was as expected for severe sepsis^12^ and higher in those with definite endocarditis (Figure 2A1) but decreased within days of commencing phage treatment. The calculated multiplicity of infection in blood (phages infused per bacterial genome detected) immediately after phage administration (MOI_input_)^13^ was generally >100. Phage DNA was detected up to 12 hours after administration and viable phages were recoverable from blood up to an hour after administration, as in previous reports^14^, for up to five days after starting therapy in our cohort (data not shown). In one patient (WMP0011), identical numbers of viable phage were retrieved from both sides of the oxygenator in an extra-corporeal membrane oxygenation (ECMO) circuit and no viable phages were detected in effluent collected 15 minutes after AB-SA01 dosing.

Clinical inflammatory markers (white cell count and C-reactive protein) declined in most patients during or soon after phage therapy commenced (Table 1). Inflammatory gene signatures in whole blood transcriptomes of definite IE patients (n=4) declined significantly within 72 hours of commencing bacteriophage therapy (Figure 2C). Analysis of gut microflora using *16S rRNA* amplicon sequencing before and after bacteriophage therapy revealed no obvious effect on population diversity overall nor evident depletion of *Firmicutes* (data not shown).

## DISCUSSION

Propensity-based/ case-matching analyses were rejected as unfeasible due to the large number of variables and the small size of our cohort and because the study was designed for an endpoint of safety rather than mortality. Nevertheless, our experience indicates that controlled trials of bacteriophage therapy are warranted to answer some of the many questions that remain, and the relatively standardised nature of antimicrobial therapy for *S. aureus* infection makes this a good model. Phage-based products have previously received U.S. FDA licensure for food safety applications and a recent study showed reduction of bacterial burden in superficial burns^15^. Here we show for the first time that that IV-administered investigational phage therapy produced under GMP conditions is safe and well-tolerated in severe sepsis.

The reticuloendothelial system is the most important factor in initial clearance of phage from the bloodstream^13^ and many successful animal studies use a single dose of phage but a few of our patients had bacteriophage detectable in blood up to 12 hours after dosing and we detected traces of phage (and bacterial) DNA in valve tissues resected after 14 days (WMP010). Most of our cohort, however, had cleared bacterial and phage DNA from blood within a week of starting phage therapy. Optimal MOI_input_ after a dose of 10^9^ PFU is likely to be adequate^13^ for any initial *in vitro* synergy between phages and antibiotics^4^ but the benefit of adding bacteriophages to optimal antibiotic therapy remains uncertain. The observed decline in inflammatory responses shortly after bacteriophage administration is consistent with previous animal data^16^ and may be important in suppressing destructive disease processes but may simply reflect 4-10 days of preceding antibiotics. Robust controlled trials are needed to determine the contribution of adjunctive phage therapy in severe *S. aureus* infections.

### Added value of this study

This study is the first of its kind where an investigational GMP-quality phage product was administered intravenously to treat severe *S. aureus* infections. Administration of AB-SA01 was followed by significant reduction in bacterial burdens and inflammatory profiles and by no adverse events. Development of resistance to AB-SA01 was not detected. Our data indicate that AB-SA01 is suitable as adjunctive therapy for severe *S. aureus* infections using the dosing strategy described, and that controlled clinical trials are an important next step.

## Acknowledgments

We would like to thank Joey Lai and Li Ma at WIMR and the staff of the Pathogen Genomics Unit at Westmead Hospital and Ramaciotti Centre (UNSW) for technical advice and sequencing support, Angela Netluch and Patricia Fa for pharmacy support, Stephanie Chan, Katherine Garnham, Sonya Natasha Hutabarat, Cecilia Li, Chris Robson, Andrew Ginn, Belinda Roychoudry, Neela Joshi Rai and the staff of the NSW Pathology Microbiology laboratory at Westmead Hospital for help with data and sample collection and Bethany Bowring for lab assistance.

## Author contributions

JI conceived the project and devised the treatment protocol. SMa, RCYL and JH were involved in amendment of the protocol v2.0 and v3.0. SMa and JH created the case report forms. JH screened patients. JI, SMa, TG, IS consulted on suitability of patients. RC was consulted for all IE patients. JI, RCYL, APF designed the experiments. JI, APF, RCYL, JH and SMa wrote the paper. AFP, RCYL, JH collected and processed samples, and analysed data. JH collected retrospective patient data. APF conducted and analysed phage-susceptibility. APF, RCYL and JI analysed bacteria and phage kinetics data. NLBZ analysed isolate WGS data. CV analysed faecal amplicon sequencing data. RCYL and APF analysed transcriptome data. WBTT participated in case finding and therapeutic management. AmpliPhi Biosciences collaborated in design of the protocol and prepared essential supportive documents (including the Pharmacy Manual and AB-SA01 Product Information and certificates of analysis), critically reviewed individual patient narratives and produced AB-SA01 investigational phage product under GMP conditions.

### Diagnoses

A: aortic valve
M: mitral valve
IE: infective endocarditis
PVE: prosthetic valve endocarditis
RIE: right-sided infective endocarditis
T: tricuspid valve
OM: osteomyelitis.

### S. aureus ST

ST: *S. aureus* sequence type (Methicillin-resistant *S. aureus* in parentheses)

* AB-SA01 EOP reduced by >2.0 log_10_ *in vitro* compared to SA-5863

### Predicted % mortality

APACHE II: Acute Physiology and Chronic Health Evaluation (% in-hospital mortality), absolute score in parentheses
CCI: Charlson Comorbidity Index (predicted % 1-year mortality)
IE: Infective Endocarditis Score (predicted % 6-month mortality)

### Antibiotics and phage therapy

^a^Principal antibiotics (used for >72hours)

CEF: Cefazolin
CIP: Ciprofloxacin
CLI: Clindamycin
FLU: Flucloxacillin
MEM: Meropenem
RIF: Rifampicin
VAN: Vancomycin
^b^ (days prior): days of effective antistaphylococcal antibiotic therapy prior to starting bacteriophage therapy.
CRP: C-reactive protein (mg/L), Reference value: ≤3mg/L
WBC: White blood cells (×10^9^/L), Reference value: ≤3mg/L
ND: not done
^c^ Pre: 24hours before phage administration
^d^ Pre: 48hours before phage administration
^e^ D14: end of phage therapy (+/-48hours)

### Comments

ECMO: Extracorporeal Membrane Oxygenation
CRRT: Continuous Renal Replacement Therapy

